# JAK1/2 Inhibition Restores Tissue Integrity by Rewiring Type 1/2 Immunity Toward Targetable IL-17 Responses

**DOI:** 10.64898/2026.10.30.763025

**Authors:** Regina Jin, Karoline Strobl, Lena Artner-Gent, Johannes Tüchler, Marina Gladkova, Ulrike Mann, Matthias Farlik, Iros Barozzi, Maria Sibilia, Thomas Bauer

## Abstract

Effective immune responses require coordinated communication between immune and epithelial cells to eliminate pathogens while limiting tissue damage. Disruption of this balance drives chronic inflammation and progressive tissue destruction. The Janus kinase (JAK) pathway coordinates cytokine signaling across these compartments and has emerged as a therapeutic target in inflammatory diseases. However, the consequences of JAK inhibition within a complex inflammatory environment remain incompletely understood.

Here, we dissect the molecular, cellular, immunological and phenotypical effects of topical JAK1/2 inhibition in a chronic model of tissue-destructive skin inflammation driven by barrier dysfunction and bacterial superinfection. Single-cell transcriptomics and functional profiling revealed that JAK1/2 blockade selectively suppressed pathogenic CD8⁺ T cells and their IFN-γ production while shifting Th2-polarized CD4⁺ T cells toward a Th17 phenotype, preserving type 3 immunity consisting of IL-17A-producing γδ-T cells and neutrophil responses. This immune rebalancing occurs rapidly and concomitantly with the restoration of epithelial barrier integrity through tight junction reinforcement, followed by recovery of antimicrobial defenses and reduced bacterial burden. Combination of JAK1/2 and IL-17A blockade further improved tissue homeostasis by targeting residual type 3-driven pathology.

Thus, JAK1/2 inhibition acts as selective immunomodulator rather than a broad immunosuppressant, targeting pathogenic type 1/2 responses while preserving type-3-associated antimicrobial defense programs. These findings reveal how JAK1/2 inhibition coordinates immune rebalancing with epithelial repair and support its therapeutic potential in chronic type 1/2-driven inflammatory disease, while identifying residual IL-17A-mediated pathology as a target for combination therapy.

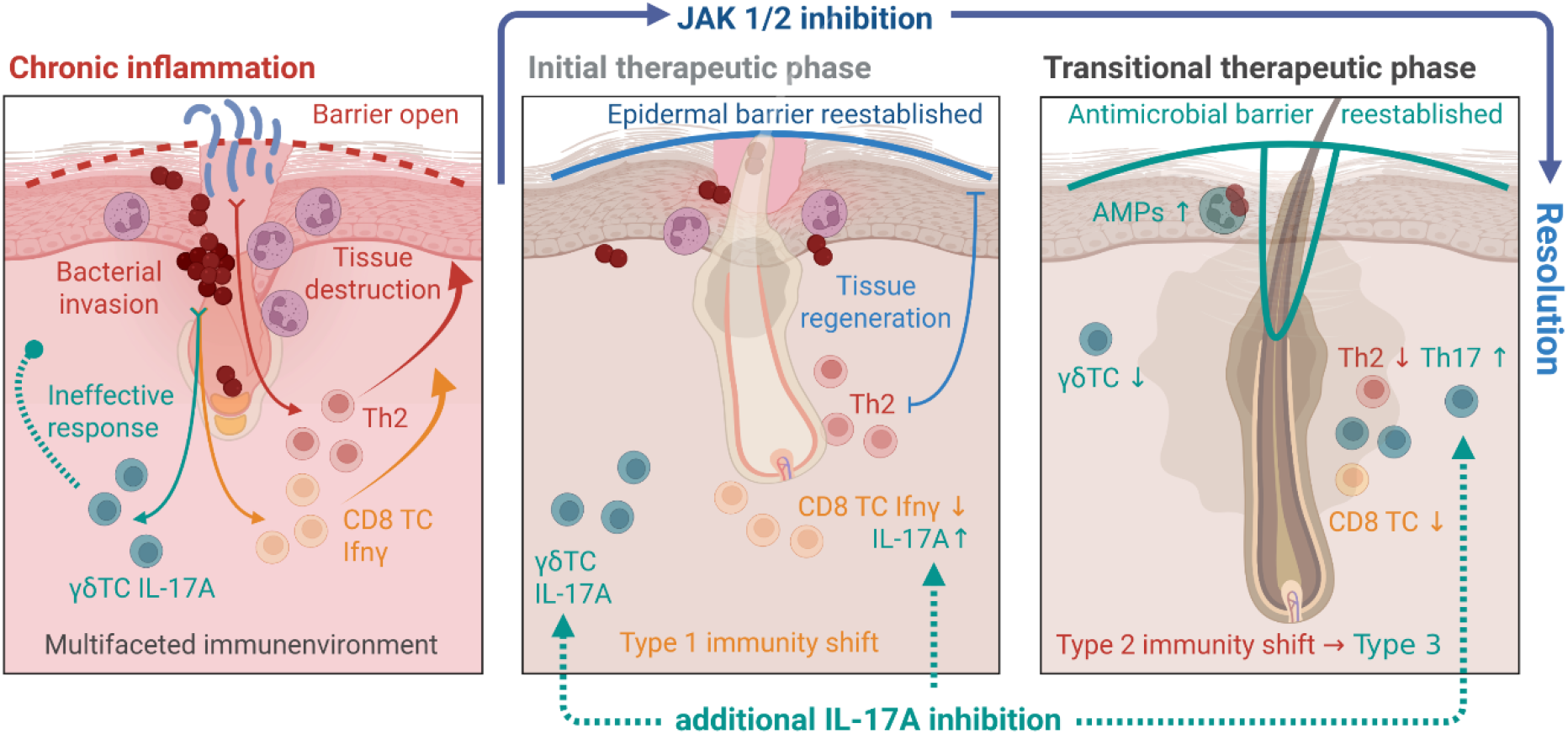
Graphical abstract.

## Introduction

Communication between cells is essential for the immune system to respond and adapt to various danger signals, coordinate effective immunity and interact with structural cells. This ensures efficient pathogen clearance while minimizing collateral tissue damage. However, when this balance is disrupted, it can result in a prolonged, unresolved inflammatory response. The resulting chronic disease leads to progressive tissue destruction, can trigger various malignancies and impairs microbial control at barrier sites.^1–3^

The Janus kinase (JAK)-signal transducer and activator of transcription (STAT) pathway is central to this process by mediating intracellular signaling transduction of extracellular immune modulators (e.g. cytokines).^4^ As such, the JAK-STAT pathway is an attractive therapeutic target. In recent years, JAK inhibitors (JAKis) have emerged as a novel drug class in the treatment of various inflammatory conditions including skin diseases.^5^ Several JAKis have received FDA approval for cutaneous conditions including oral baricitinib (JAK1/2 inhibitor) for atopic dermatitis (AD) and alopecia areata (AA), topical ruxolitinib (JAK1/2 inhibitor) for AD and vitiligo, and deucravacitinib (TYK2 inhibitor) for psoriasis.^6^ Additionally, JAKis have also shown promise in difficult-to-treat conditions such as scarring alopecia and toxic epidermal necrolysis.^7–10^

Despite the growing popularity of JAKis, our limited immunological understanding of the impact of different JAKis prevents us from predicting their spectrum of targets and hinders their use as powerful precision medicine tools. The four JAK proteins JAK1, JAK2, JAK3 and TYK2 are associated with different cytokine receptors and in different combinations are responsible for the signal transduction of over 50 different cytokines.^5^ Thus, one single JAKi has the potential to act on a broad range of immunological effects simultaneously. This mechanism of action may be especially valuable in inflammatory skin conditions that involve more than one or two key causal cytokines, where specific monoclonal blocking antibodies are insufficient.^11^ On the other hand, this broad suppression of the immune system may also cause unwanted and unexpected adverse events including increased infection risk.^12^ Therefore, it is crucial to know which aspects of a multifaceted immune response are specifically affected and how.

We recently identified the JAK-STAT pathway as a therapeutic target in scarring hair follicle destruction and skin inflammation induced by EGFR-deficiency.^7^ In a mouse model with hair follicle-specific EGFR deletion (EGFR^ΔEgr^^2^), a 4-week treatment regimen with a topical JAK1/2 inhibitor ameliorated skin inflammation, an effect accompanied by the partial recovery of hair growth.^7^ In humans, the potential of JAKi therapy in patients with EGFR inhibitor induced skin rash was also demonstrated recently in a pilot clinical trial.^13^

Mechanistically, EGFR deficiency disrupts its downstream SOS-ERK cascade which results in barrier failure during hair eruption, inducing a type 2-dominated immune response including robust TSLP production.^14,15^ The compromised barrier simultaneously facilitates the entry of commensal microbiota through the follicular opening, provoking additional type 1 and type 3 responses. The resulting multifaceted inflammatory environment favors *Staphylococcus aureus* (*S. aureus*) dysbiosis. Taken together, the specific inflammatory signature, barrier disruption, tissue destruction and antimicrobial dysfunction are all hallmarks of AD.^16,14,17^

Here, we used this preclinical model of chronic skin disease to investigate the targeted therapeutic effects of JAK1/2 inhibition at the organismal, immunological, single-cell, and molecular levels. Our findings demonstrate that JAK1/2 inhibition restores epidermal and hair follicle barrier integrity, reverses tissue destruction, and resolves persistent superinfection by selectively dampening type 1 and type 2 effector responses while rewiring immunity to antimicrobial type 3 immune programs. Combinatorial blockade of JAK1/2 and IL-17A further enhanced therapeutic efficacy. These results position JAK inhibitors as precise immunomodulatory agents rather than broadly immunosuppressive drugs, supporting their targeted use in chronic inflammatory diseases within precision medicine frameworks.

## Results

### Prolonged EGFR dysfunction drives chronic skin inflammation with a multifaceted inflammatory environment in mice and humans

EGFR signaling is essential for maintaining cutaneous homeostasis and barrier integrity.^18,19^ Although prolonged EGFR dysfunction induces inflammatory skin pathology, the immunological mechanisms linking epithelial dysfunction to chronic tissue-destructive inflammation remain incompletely understood. To investigate how sustained EGFR dysfunction shapes the inflammatory environment, we analyzed gene expression profiles of scalp biopsies of patients receiving long-term EGFR inhibitor treatment^20^ and those from back skin of adult mice lacking the EGFR in the hair follicle compartment (EGFR^ΔEgr2^).^7^

Semantic similarity analysis of enriched Gene Ontology biological processes in human long-term EGFRi-treated skin revealed a multifactorial immunological environment including chemokine, T-cell activation, type II interferon production and active antimicrobial defense alongside processes associated with tissue disruption (Fig. 1A). Consistent with this broad inflammatory signature, analysis of candidate immune-regulatory genes revealed dysregulation of mediators associated with type 1 (e.g. *CXCL10*), type 2 (e.g. *IL4I1*) and type 3 (e.g. *IL17A*) immune responses (Fig. 1B and Fig. S1A and B).

**Fig. 1.**
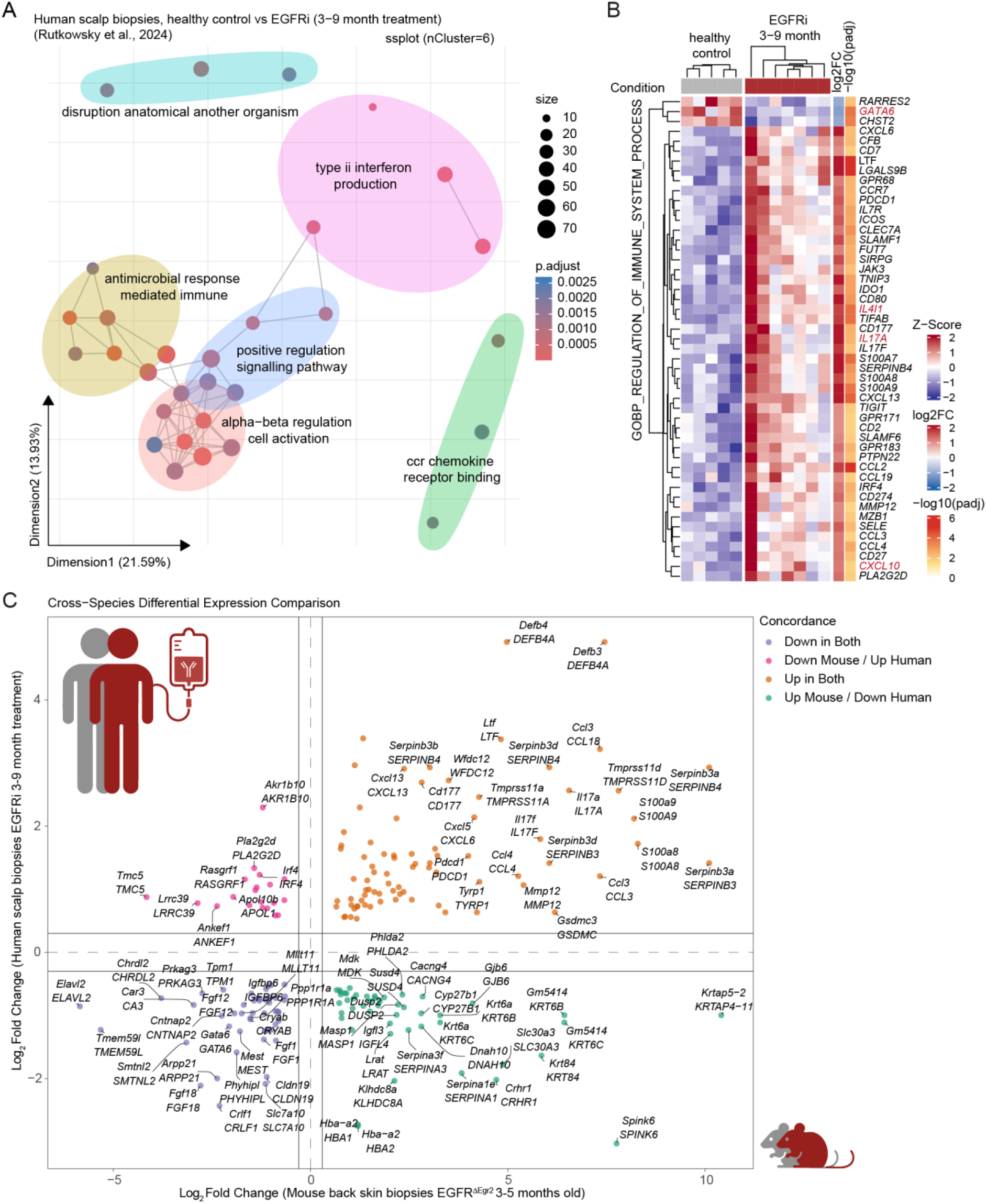
Human skin inflammation during long-term EGFR inhibitor treatment recapitulates multifaceted inflammatory signatures in aged EGFR-deficient mice. RNA sequencing data of scalp biopsies of patients treated 3-9 months with EGFRi (n = 7) and healthy controls (HC, n = 5) from Rutkowski *et al.*^20^ were re-analysed and compared to mRNA sequencing data of full-thickness back skin of EGFR^ΔEgr2^ mice and WT littermates (3-5 months old). **(A)** Semantic Space plot of enriched gene sets in EGFRi-treated patients determined by gene set enrichment analysis (GSEA) using the Gene Ontology collection. The Semantic Space plot clusters similar gene sets. **(B)** Heatmap of significantly differentially expressed genes between EGFRi-treated and HC (adjusted p-value < 0.05 & absolute log2 fold change ≥ 1) which overlap with the Gene Ontology gene set GO:0002682. **(C)** Comparison of significantly differentially expressed genes (adjusted p-value < 0.05 & absolute log2 fold change ≥ 0.5) in EGFRi-treated patients and their mouse orthologs in EGFR^ΔEgr2^ mice.

To assess whether this inflammatory environment is conserved across species, we compared differential gene expression between the human EGFRi-treated skin and back skin from 3-5-month-old EGFR^ΔEgr2^ mice. The mouse transcriptome showed substantial overlap with the human inflammatory signature, supporting the conservation of key inflammatory features across species (Figure 1C). Notably, both datasets exhibited reduced *Gata6* expression, consistent with the dysfunction of upper hair follicle cells.

Together, these findings indicate that chronic EGFR dysfunction is associated with a species-conserved, multifaceted inflammatory environment involving type 1, type 2, and type 3 immune responses, accompanied by epithelial dysfunction and tissue disruption.

### Hallmarks of the transitional therapeutic stage of JAK1/2 inhibition in chronic inflammation

We could previously show that JAK1/2 inhibition in bald EGFR^Δep^ (Keratin Krt5-epidermal-specific EGFR deletion) and EGFR^ΔEgr2^ (EGR2-hair-follicle-specific EGFR deletion) mice leads to beneficial therapeutic effects including visible hair regrowth after 1 month of treatment^7^ (defined as resolution phase, Fig. S2A).

To investigate the molecular mechanism of topical JAK1/2 inhibition on each epidermal and immunological cell type during the transition from multifaceted inflammation to resolution, we performed single-cell RNA sequencing (scRNA-seq) after 2 weeks of treatment (defined as the transitional therapeutic phase, Fig. 2A and S2A). Adult EGFR^ΔEgr2^ mice with visible hair loss and an open skin barrier together with wild-type (WT) mice were topically treated with a JAK1/2 inhibitor (JAK1/2i, 3% ruxolitinib in DMSO) or vehicle (DMSO). On the organism level, therapy success was already noticeable at this timepoint by the visible darkening of the back skin due to the reappearance of melanin-rich hair shafts in the treated mice as compared to the control group (Fig. 2B). The epidermal barrier function of the JAK1/2i-treated EGFR^ΔEgr2^ mice was also improved as measured by reduced transepidermal water loss (TEWL, Fig. 2C).

**Fig. 2.**
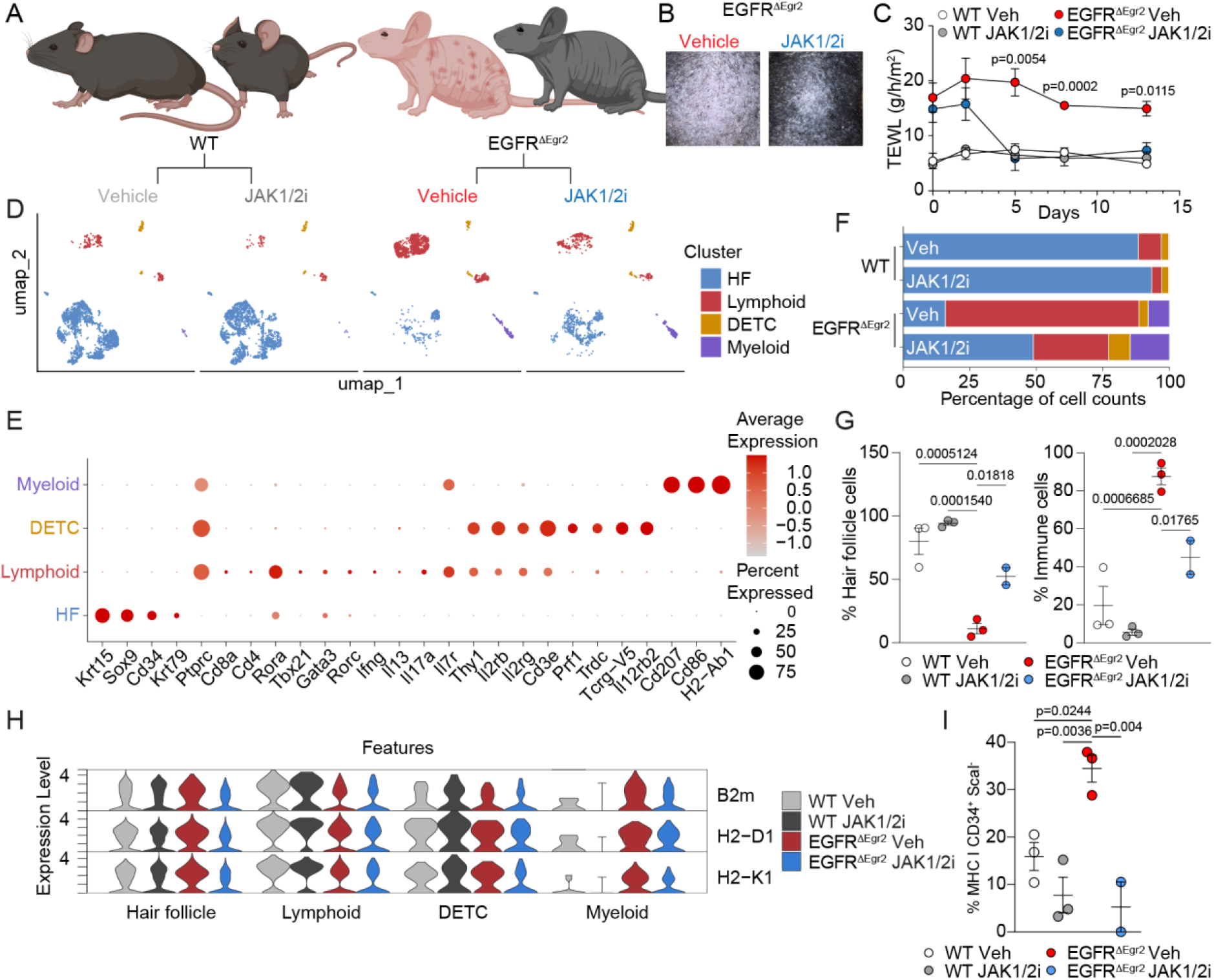
Single-cell profiling reveals the epidermal immune and hair follicle cell landscape following topical JAK1/2 inhibition. EGFR^ΔEgr2^ mice and WT littermates were treated topically with the JAK1/2i ruxolitinib for 2 weeks. Immune cells and hair follicle cells were then isolated from the epidermis and subjected to single-cell RNA sequencing. **(A)** Schematic of the experimental conditions. **(B)** Representative pictures of mouse backskin after 2 weeks treatment. **(C)** Transepidermal water loss (TEWL) measured at indicated timepoints (n ≥ 2) showing mean ± standard error of the mean (SEM). P-values (EGFR^ΔEgr2^ Ctrl vs. JAK1/2i) were determined by one-way ANOVA with Tukey’s multiple comparisons test. **(D)** Uniform manifold approximation projection (UMAP) of main classes detected in the scRNA-seq. Dots represent single cells and are colored by assigned main class. **(E)** Expression of selected cell type marker genes. **(F)** Cellular composition of main clusters across conditions. **(G)** HF or immune cell abundance in combined immune/HF gate in epidermis of backskin measured by flow cytometry. **(H)** Expression of class I antigen presentation genes in main classes of the scRNA-seq data. **(I)** Percentage of MHC-I+ hair follicle stem cells quantified by flow cytometry. Data in (G) and (I) are shown as mean ± SEM with each dot representing one mouse. P-values are shown above the relevant groups and were determined by one-way ANOVA with Tukey’s multiple comparisons test.

Having confirmed therapeutic efficacy superficially, we next analyzed the scRNA-seq data of FACS-sorted single CD45^+^ immune cells and CD45^-^Sca1^-^hair follicle (HF) cells from the epidermal compartment of these mice (see Fig. S2B for gating strategy). After quality control steps, each of the four conditions featured at least 1,259 single cells from at least two individual mice per group. The cells were divided into four main classes based on known marker gene expression: HF, Lymphoid, dendritic epidermal T cells (DETC), and Myeloid (Fig. 2D, 2E and S2C). The HF class is characterized by bulge markers *Krt15*, *Sox9*, and *Cd34* in addition to the upper HF (uHF) marker *Krt79* (Fig. 2E). The Lymphoid class consisted of a mixed population of epidermis-infiltrating lymphocytes including T cells and innate lymphoid cells (ILCs). These were characterized by expression of *Rora*, *Tbx21*, *Gata3*, *Rorc*, and *IL7r* (Fig. 2E). The resident DETCs could be distinguished by *Tcrg-V5* (Fig. 2E and S2C). The Myeloid class comprised mostly Langerhans cells (LC), expressing Langerin (*Cd207*) and MHC-II genes, and infiltrating neutrophils (Fig. 2E). In the vehicle-treated EGFR^ΔEgr2^ group, the infiltrating Lymphoid cluster dominated the cellular composition while the HF compartment was strongly reduced compared to both WT conditions, reflecting the already advanced inflammatory hair follicle destruction (Fig. 2F). JAK1/2 inhibition in the EGFR^ΔEgr2^ group resulted in a re-emergence of the hair follicle compartment and a reduction of infiltrating lymphocytes in line with the onset of the therapeutic effect during this transitional phase (Fig. 2F). WT mice treated with JAK1/2i exhibited no major changes in barrier integrity, and only slight decreases in the proportion of Lymphoid cells (Fig. 2C, D and F). These quantitative cellular changes in HF and immune cells were confirmed by additional flow cytometry analysis of the same mice used for the scRNA-seq (Fig. 2G). JAK1/2 inhibition also down-modulated the expression of the class I antigen presentation machinery in the HF compartment, restoring the immune privilege of the HF stem cells (Fig. 2H and 2I), as we have previously shown for the resolution phase.^7^

In summary, 14-day topical JAK1/2i treatment initiated hair re-growth, normalized cutaneous barrier function, reduced epidermis-infiltrating lymphocytes and reduced inflammation, defining the transitional phase of the therapy in our preclinical model.

### JAK1/2 blockade restores hair follicle architecture and reconstitutes the upper follicular barrier integrity

To characterize the rudimentary HF compartment in bald EGFR^ΔEgr2^ mice and assess the cell-type-specific effects of JAK1/2i, we next performed targeted sub-clustering analysis of the HF populations. Based on known marker gene expression, the cells could be assigned to two distinct upper HF (uHF) clusters (uHF_Lrig1: *Lrig1* and uHF_AMP (antimicrobial peptide): *Defb6*), one cluster comprising sebaceous gland (SG: *Pparg*), one isthmus cluster (*Lgr6, Nrtn*), one inner bulge cluster (IB: *Fgf18*), four outer bulge clusters (OB: *Cd34*, *Krt24*, *Postn*) and a heterogeneous Non_KC cluster expressing melanocyte (*Dct*) and fibroblast markers (*Vim*) (Fig. 3A and S3A-C). Compared to WT mice, EGFR^ΔEgr2^ mice had nearly undetectable IB and uHF clusters (Fig. 3A and 3B). Isthmus and the outer bulge clusters OB_Postn, OB_Krt24, and OB_Cdk8 were strongly reduced, indicating the ongoing destruction of the stem cell compartment (Fig. 3A and 3B). Most remaining cells in EGFR^ΔEgr2^ mice were associated with the OB_Camk1d and SG clusters (Fig. 3B). Cells from the OB_Camk1d cluster expressed markers typical of the outer bulge, but at lower levels than the other outer bulge clusters (Fig. S3A). JAK1/2i treatment led to an expansion of all HF populations in relation to total cells (Fig. 3B). We then performed RNA velocity analysis to predict which remaining hair follicle stem cell populations might contribute to the recovery of the hair follicle. JAK1/2i restored the transcriptional dynamics of the niche (Fig. 3C). Within the JAK1/2i-treated EGFR^ΔEgr2^ HF compartment the RNA velocity streamlines appeared more directional and pointed from the OB_Camk1d cluster towards the other outer bulge populations (Fig. 3C). We also noted that the OB_Camk1d cluster of EGFR^ΔEgr2^ mice displayed a drop in unspliced-to-spliced ratio, which is restored with JAK1/2i, suggesting a transcriptional reactivation of the niche (Fig. 3D and S3D). The uHF clusters re-emerged during JAKi treatment but did not have any obvious trajectories to or from bulge clusters (Fig. 3C).

**Fig. 3.**
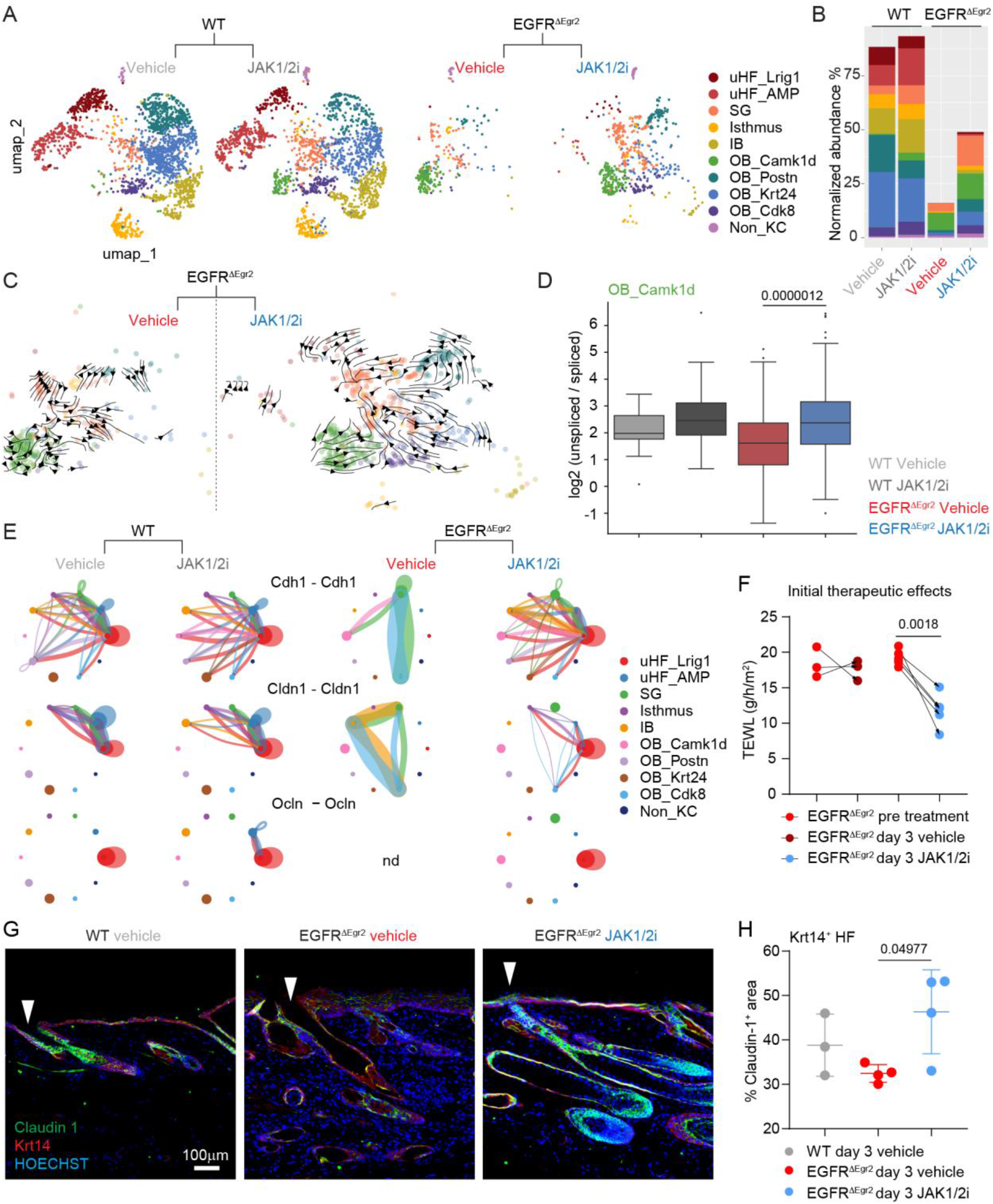
JAK1/2 inhibition restores barrier function through restoration of the upper hair follicle. HF class from scRNA-seq was subsetted and reclustered to resolve subclusters. **(A)** UMAP of HF subclusters with dots colored by expression of selected cluster marker genes and with each condition shown separately. uHF = upper hair follicle, SG = sebaceous gland, IB = inner bulge, OB = outer bulge, Non_KC = Non keratinocytes including fibroblastic cells and melanocytes. **(B)** Cellular abundance normalized to all cells (including immune cells) within the condition. **(C)** RNA velocities of EGFR^ΔEgr2^ conditions projected onto UMAP embeddings and visualized as streamlines. **(D)** Log₂ unspliced/spliced transcript ratio in the OB_Camk1d cluster across conditions, determined from velocyto spliced/unspliced counts. Boxes show the median and interquartile range across cells. q-values are from two-sided Mann–Whitney U tests of pairwise differences between conditions (see Methods). Only q-values below 0.05 are shown. **(E)** Circle plot depicting Cdh1 – Cdh1, Cldn1 – Cldn1 and Ocln – Ocln cell-cell interactions as determined by CellChat. Vertices (dots depicting clusters) are scaled proportionally to cluster size, while edges (connections between clusters) are weighted according to the number of predicted interactions and colored by sender. **(F)** TEWL values measured pre-treatment and after 3 days of JAK1/2i or vehicle (n ≥ 3). P-values are shown if ≤ 0.05 and were determined by 2-way ANOVA with Sidak’s multiple comparisons test. **(G)** Representative images of claudin-1 and keratin-14 immunofluorescence staining of back skin sections after 3 days with JAK1/2i. **(H)** Quantification of claudin-1^+^ area in hair follicle region (n ≥ 3). Data are shown as mean ± standard deviation (SD). P-values are shown if ≤ 0.05 and were determined by 1-way ANOVA followed by Tukey’s multiple comparisons test.

We next used CellChat to infer cell-cell interactions and communication between the different HF clusters (Fig. S3E). In line with the barrier defects, deregulated interactions associated with cell-to-cell adhesion were among the most prominent. The tight junction proteins claudin-1 (*Cldn1)* and occludin (*Ocln)* were mainly detected in the uHF clusters in the WT HF (Fig. 3E). Similarly, E-cadherin (*Cdh1)* interactions, which form adherens junctions, were detected in the uHF_Lrig1 cluster (Fig. 3E). These interactions were either undetectable or emerged between different clusters in the vehicle-treated EGFR^ΔEgr2^. Importantly, *Cldn1*, *Ocln* and *Cdh1* interactions from the uHF_Lrig1 cluster were re-established in the JAK1/2i-treated EGFR^ΔEgr2^ (Fig. 3E). This prompted us to check for the earliest timepoint of barrier restoration. Interestingly, epidermal water evaporation was already normalized during the initial therapeutic phase as early as three days into the JAK1/2i treatment (Fig. 3F). Concomitantly, claudin-1 immunostaining re-emerged in the upper hair follicle (Fig. 3G and 3H).

In conclusion, JAK1/2i treatment in EGFR^ΔEgr2^ mice led to the recovery of diverse HF subpopulations, most importantly, the upper HF populations of the hair canal, which express anti-microbial peptides and tight junction and adherens junction proteins important for the protective cutaneous barrier function.

### JAK1/2 inhibition specifically targets type 1 and 2 immunity, while preserving and rewiring to type 3 responses

In order to assess the effect of JAK1/2i on the different immune cell subsets, we next re-clustered the Lymphoid compartment from the main dataset (see Fig. 2D). The bulk of the immune infiltrate consisted of γδ (*Trdc*), CD4, CD8 and regulatory T cells (Tregs, *Foxp3*, Fig. 4A and 4B). The γδ T cells (γδTC) encompass two subsets, resulting in a total of 5 Lymphoid clusters annotated according to the expression of known cell-type-specific marker genes (Fig. 4A, S4A and S4B). ILCs grouped together with their respective T cell counterparts according to immune polarization. The CD4 cluster featured markers of CD4 T cells and ILC2s reflecting type 2 immunity, while the CD8 cluster expressed markers of CD8 T cells, NK cells and ILC1s reflecting type 1 immunity (Fig. S4A). The two γδTC clusters included the ILC3s and represented type 3 immunity. Lastly, Tregs formed a cluster of their own with some cells included in the main CD4 cluster.

**Fig. 4.**
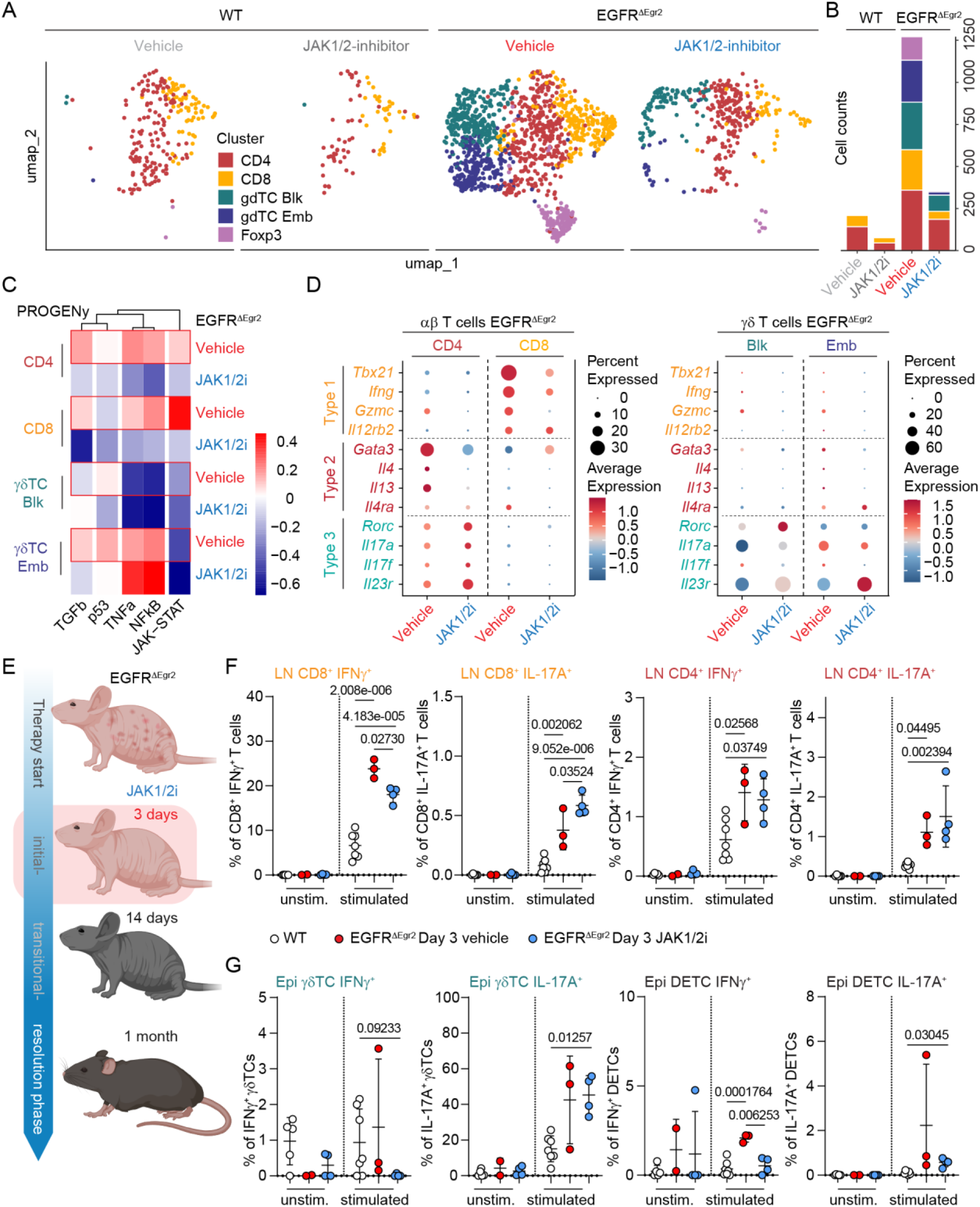
JAK1/2 inhibition differentially modulates T cell responses and reveals Th17 reprogramming. Lymphoid compartment from the scRNA-seq of EGFR^ΔEgr2^ mice and WT littermates treated with JAK inhibitor or vehicle was subsetted and re-analysed on its own. **(A)** Lymphoid subclusters visualized as UMAP split by condition. **(B)** Cellular abundance of each cluster in each condition depicted as percentage of all cells in each condition. **(C)** Heatmap of PROGENy activity scores for depicted pathways. **(D)** Expression of T helper genes in T cell clusters from vehicle– and JAK1/2i-treated EGFR^ΔEgr2^ mice. **(E)** Scheme depicting the therapeutic phases of JAK1/2i treatment. **(F-G)** EGFR^ΔEgr2^ and WT mice were treated topically with JAK1/2 inhibitor or vehicle for 3 days. Cells from skin-draining LNs or backskin epidermis were isolated and incubated *ex vivo* with or without ionomycin and phorbol 12-myristate 13-acetate, followed by intracellular cytokine staining. Unstimulated and stimulated conditions are shown in the plots. **(F)** Flow cytometry analysis of LN T cells showing percentages of cells expressing IFNγ or IL-17A (n ≥ 3). Data are shown as mean ± SD. P-values were determined by one-way ANOVA with Tukey’s multiple comparisons test. **(G)** Flow cytometry analysis of backskin T cells showing percentages of cells expressing IFNγ or IL-17A (n ≥ 3). Data are shown as mean ± SD. P-values were determined by Brown-Forsythe and Welch ANOVA tests to account for unequal variances between groups, followed by Games-Howell’s post hoc test.

As expected, the WT conditions had very few Lymphoid cells in total, which were mainly from CD4 and CD8 clusters (Fig. 4B). The inflammatory environment of EGFR^ΔEgr2^ mice was represented by dramatically expanded CD4 and CD8 compartments along with additional γδTC and Treg infiltrates (Fig. 4B). Transitional JAK1/2i treatment of EGFR^ΔEgr2^ mice led to a decrease in all Lymphoid clusters compared to vehicle-treated EGFR^ΔEgr2^ mice (Fig. 4B).

Interestingly, the PROGENy score for JAK-STAT pathway activity was highest in the CD8 and CD4 compartments of the vehicle-treated EGFR^ΔEgr2^ mice as compared to the γδTC compartments (Fig. 4C). Concomitant with that observation, the JAK-STAT pathway along with the pro-inflammatory TNFα and NF-κB pathways were effectively dampened in these clusters under JAK1/2i therapy. Strikingly, the type 3-polarized γδTC clusters had relatively low JAK-STAT pathway activity (Fig. 4C). The gdTC_Emb cluster, however, displayed a strong TNFα and NF-κB activity, which was not inhibited, but rather enhanced by the JAK1/2i (Fig. 4C).

We next examined the immune polarization status during the transitional phase of the JAK1/2i therapy. JAK1/2i-treatment in EGFR^ΔEgr2^ mice specifically targeted the expression of type 1 and type 2-associated genes in the CD4 and CD8 clusters, respectively (Fig. 4D). Surprisingly, unlike the strong reduction of cytotoxic responses in the CD8 compartment, the CD4 compartment shifted their polarization from Th2 to Th17 (i.e. *Gata3*, *Il4, Il4ra* and *Il13* were downregulated, while *Rorc, Il17a, IL17f* and *Il23r* were upregulated; Fig. 4D, left panel, and S4C). Supporting the weak JAK-STAT activity in the two γδTC compartments, their signature gene expression did not change dramatically with selected genes even elevated (i.e. *Rorc* and *Il23r*; Fig. 4D, right panel). In contrast to the recruited γδTCs, the skin-resident DETCs skewed towards a type 1 phenotype, expressing the IL-12 receptor subunit gene *Il12rb2,* which was downmodulated by JAK1/2i treatment (Fig. S4C and S4D). Interestingly, they upregulated the EGFR ligand and wound healing promoter *Areg* during inflammation and downregulated it during JAK1/2i treatment (Fig. S4D).

To investigate the immediate changes, we next analyzed the initial therapeutic phase after 3 days of topical JAK1/2i treatment (Fig. 4E for schematic experimental setup). Using intracellular protein staining combined with flow cytometry, we quantified the cytokine production capacity of T cells isolated from lymph nodes (LNs) from vehicle-or JAK1/2i-treated EGFR^ΔEgr2^ mice. Confirming our scRNA-seq results, T cells from vehicle-treated EGFR^ΔEgr2^ mice produced significantly more IFNγ or IL-17A than cells isolated from WT control mice (Fig. 4F and S4E). Importantly, in the initial therapeutic phase, production of IFNγ from CD8 T cells was blunted in JAK1/2i-treated EGFR^ΔEgr2^ mice, while, unexpectantly, IL-17A production increased (Fig. 4F, left panels). The CD4 T cells, on the other hand, were not significantly affected by the initial JAK1/2i treatment (Fig. 4F, right panels). A shift from IFNγ toward IL-17A expression could also be observed in γδTCs and DETCs isolated from the epidermis of JAK1/2i-treated EGFR^ΔEgr2^ mice (Fig. 4G). The γδTCs of JAK1/2i-treated EGFR^ΔEgr2^ mice ceased residual IFNγ production while displaying robust IL-17A output (Fig. 4G, first and second panel). A similar switch could be observed in the local DETCs (Fig. 4G, third and fourth panel).

In conclusion, JAK1/2i initially shifted the CD8 compartment from type 1 to type 3 immunity, and later converted the CD4 helper compartment from Th2 to Th17 during the transitional stage. In parallel, the γδTC compartment continued to fulfill their role as main type 3 effectors during both initial and transitional stages of the therapy, while shutting down their rudimentary IFNγ production during JAK1/2i therapy.

### JAK1/2 therapy modulates neutrophilic properties and reduces bacterial burden

Taken the preserved Th17 response, we next addressed the effects of the rewired immune environment down to the myeloid compartment. It formed three subclusters, which could be assigned as LCs (*Cd207, H2-Ab1*), proliferating LCs (additional *Mki67*, histone genes) and neutrophils (*S100a8, Csf3r*, Fig. S5A-D). Interestingly, the proliferating LC subset could only be found in vehicle-treated EGFR^ΔEgr2^ mice, while neutrophils were detected in both EGFR^ΔEgr2^ conditions independent of therapeutic treatment (Fig. 5A). Immune activation markers in LCs from JAK1/2i-treated EGFR^ΔEgr2^ mice showed a mixed treatment response (Fig. S5E). For example, downregulation of *Tnfaip2*, which encodes TNFα-induced protein 2, reflects the less pro-inflammatory environment in JAK1/2i-treated EGFR^ΔEgr2^ mice.

**Fig. 5.**
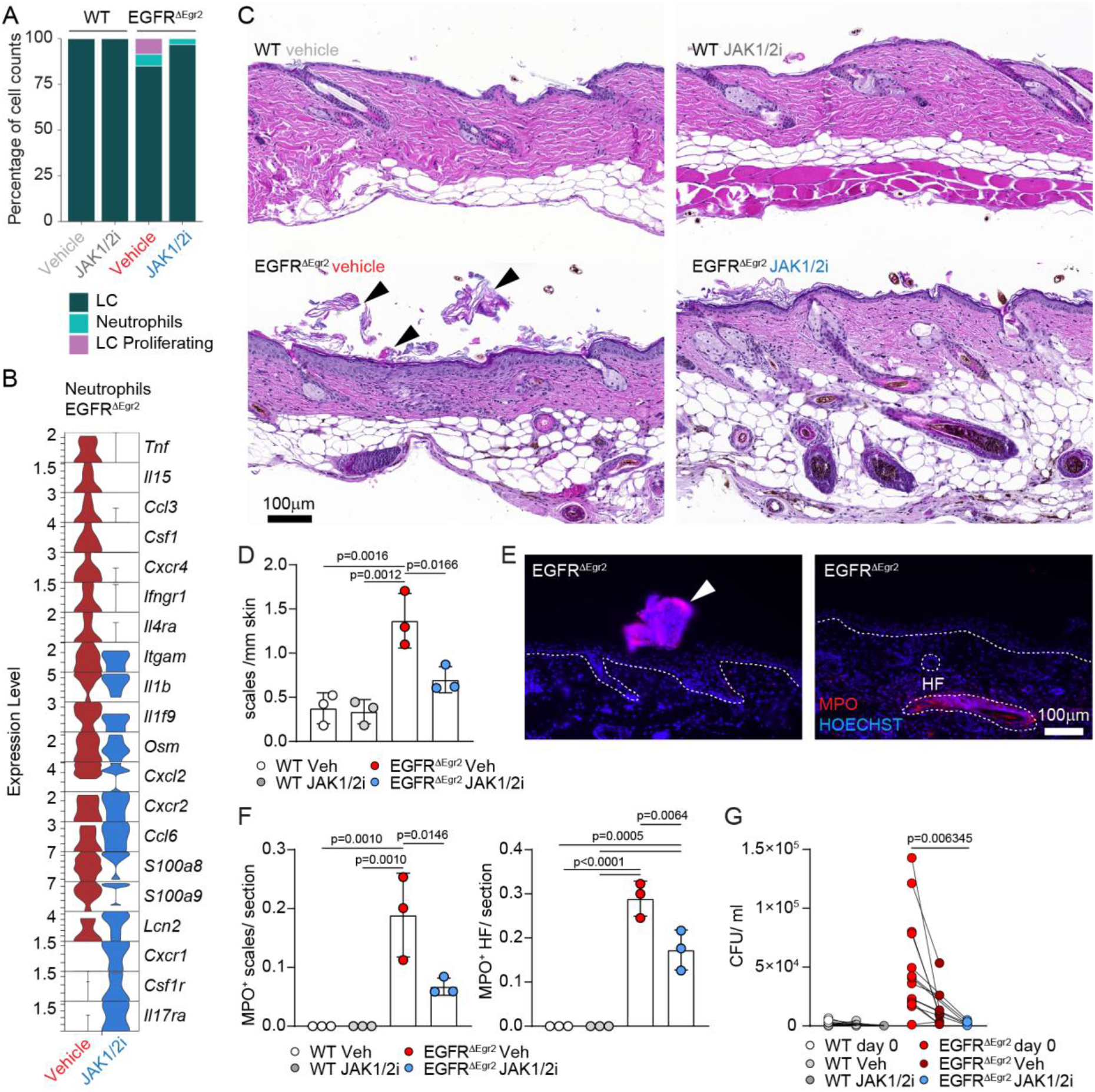
JAK1/2 inhibition restores neutrophil antimicrobial responses and promotes bacterial clearance. **(A)** Percentage of cells from each Myeloid class cluster for each group. **(B)** Violin plot depicting relative expression of selected genes in neutrophil subset of EGFR^ΔEgr2^ mice treated with vehicle or JAK1/2i for 2 weeks. **(C)** Representative images from H&E stainings of back skin from vehicle– or JAK1/2i-treated WT and EGFR^ΔEgr2^ mice. Arrows point to pustules. **(D)** Quantification of epidermal scales from backskin sections from vehicle– or JAK1/2i-treated WT and EGFR^ΔEgr2^ mice (n ≥ 3). **(E)** Representative images of immunofluorescence staining of myeloperoxidase (MPO) in backskin sections from vehicle-treated EGFR^ΔEgr2^ mice. Arrows highlight MPO-positive scales. **(F)** Quantification of MPO^+^ epidermal scales and MPO^+^ hair follicles from back skin sections from vehicle– or JAK1/2i-treated WT and EGFR^ΔEgr2^ mice (n ≥ 3). For (D) and (F), data are shown as mean ± SD and p-values were determined using a one-way ANOVA followed by Tukey’s multiple comparisons test. **(G)** Bacterial swabs were taken from the backskin of WT and EGFR^ΔEgr2^ mice before and after 2-4 weeks JAK1/2i treatment. Swabs were incubated in PBS, which was plated on Columbia agar with 5% sheep blood. Plot shows colony-forming units (CFU)/ml (n ≥ 5). P-values were calculated using a mixed-effects analysis followed by Sidak’s multiple comparisons test.

The IL-17-neutrophil axis is essential for anti-*S. aureus* defense in mice, prompting us to assess key genes for neutrophil function.^21,22^ Interestingly, Neutrophils in JAK1/2i-treated EGFR^ΔEgr2^ mice downregulated several pro-inflammatory mediators (*Tnf, Il15, Ccl3, Csf1*) along with type 1 and 2 cytokine receptors (i.e. *Ifngr1, Il4ra*, Fig. 5B). Furthermore, *Cxcr4,* a marker for senescent neutrophils that are destined for clearance^23^, is downregulated in neutrophils of JAK1/2i-treated EGFR^ΔEgr2^ mice. On the other hand, they upregulated chemokine receptors *Cxcr1* and *Cxcr2*, which bind to several CXCL chemokines including CXCL2 and CXCL6 that attract neutrophils to sites of inflammation. Importantly, we observed heightened *Il17ra* levels, together with increased expression of the antimicrobial peptides *S100a8/9* and *Lcn2* by neutrophils in JAK1/2i-treated EGFR^ΔEgr2^ mice indicating an enhanced antimicrobial defense state (Fig. 5B).

In contrast to WT mice, vehicle-treated EGFR^ΔEgr2^ mice presented with an abundant epidermal scale formation (Fig. 5C and D). JAK1/2i-treated EGFR^ΔEgr2^ mice showed significant improvement in this cutaneous phenotype. These epidermal scales and some hair follicles in EGFR^ΔEgr2^ mice were associated with myeloperoxidase (MPO)-expressing neutrophils (Fig. 5E and F). Importantly, the number of MPO^+^ scales and MPO^+^ HFs was significantly reduced after JAK1/2i treatment (Fig. 5F). Clinically most relevant, however, was the observed reduction in bacterial load on the skin of EGFR^ΔEgr2^ mice after therapeutic JAK1/2i treatment (Fig. 5G). The reduced bacterial presence aligns with the overall reduced inflammation and HF recovery. This is also evidenced by an increase in anagen stage hair bulbs in the back skin of JAK1/2i-treated EGFR^ΔEgr2^ mice compared to vehicle-treated EGFR^ΔEgr2^ mice (Fig. S5F).

In summary, the restructured immune-environment from type 1 and 2 immunity toward type 3 responses was reflected down to the neutrophil compartment adopting an IL-17-responsive anti-microbial state, accompanied by reduced epidermal scale formation and diminished bacterial burden enabling tissue regeneration.

### Rewired type 3 immunity drives IL-17A-dependent pathology

To assess the pathological aspect of the residual type 3 immunity, we next blocked IL-17A using an *in vivo* blocking antibody in combination with topical JAK1/2 inhibition for a duration of 2 weeks (Fig. 6A for schematic experimental setup). Combined JAK1/2 and IL-17A blockade maintained the beneficial effects of JAK1/2 inhibition on skin barrier integrity, as reflected by a comparable reduction in TEWL (Fig. 6B). Similar to the restored barrier function, the immune-infiltrate, including αβT-cells were also down-modulated in both therapeutic conditions (Fig. 6C, first two graphs). Interestingly, while additional IL-17A blockade did not further reduce KC-specific MHC-I expression, it resulted in a more pronounced reduction in MHC-II (Fig. 6C, last two graphs). This differential response prompted us to perform RNA-seq analysis to further characterize the molecular changes associated with combined JAK1/2 and IL-17A blockade. Most identified DEGs were downregulated under both therapeutic conditions, with the combination therapy resulting in nearly twice as many downregulated genes as JAK1/2 inhibition alone (Fig. 6D). The plethora of these genes are responsible for immunological effects, like antimicrobial responses, chemokine receptors and their G protein-coupled receptor (GPCR) signaling and neutrophil function (Fig. 6E). To determine how the different treatment regimens affected the inflammatory environment, we next generated a heatmap of the positively-correlated genes previously identified in Figure 1 by cross-species comparison of our EGFR-deficient mouse skin and human scalp skin from patients receiving long-term EGFR-inhibitor treatment (Fig. 6F). Strikingly, combined JAK1/2 inhibition and IL-17A blockade further downregulated inflammatory genes, including members of the S100A and serpin families, beyond the effects of JAK1/2 inhibition alone, restoring their expression to levels comparable to the homeostatic WT situation (Fig. 6F and Fig. S6).

**Fig. 6.**
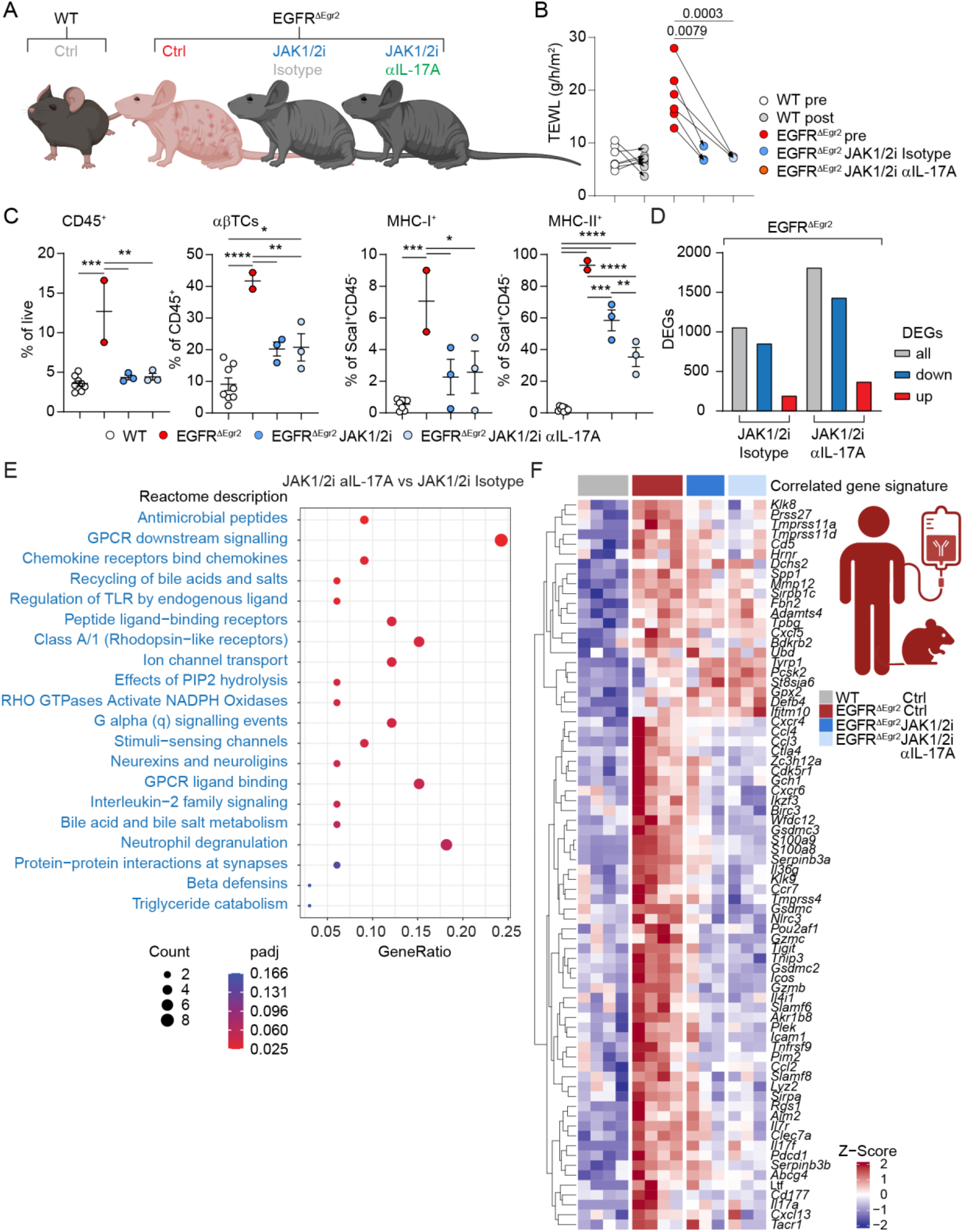
Combination of JAK1/2 and IL-17A blockade further reduces inflammation while preserving JAK1/2 inhibitor-mediated barrier restoration. EGFR^ΔEgr2^ mice were treated with JAK1/2i and concomitantly with isotype control or anti-IL-17A blocking antibody for two weeks. **(A)** Scheme of experimental groups. **(B)** TEWL before and after treatment (n ≥ 3). P-values were determined with a two-way ANOVA followed by Sidak’s multiple comparisons test. **(C)** Epidermal cells were isolated from ear skin and underwent FACS. Dotplots showing percentages of CD45^+^ immune cells within all live cells, αβ T cells within CD45+ cells, and MHC-I^+^ or MHC-II^+^ cells within the IFE compartment. Data are shown as mean ± SD. P-values are shown above the relevant groups (n ≥ 3) and were determined by one-way ANOVA with Tukey’s multiple comparisons test. **(D)** Back skin biopsies underwent mRNA sequencing analysis (n ≥ 3). Number of differentially expressed genes (DEGs, p-value ≤ 0.05 & absolute log2 fold change ≥ 1.0) in EGFR^ΔEgr2^ JAK1/2i or EGFR^ΔEgr2^ JAK1/2i anti-IL-17A conditions in comparison to control EGFR^ΔEgr2^. **(E)** Downregulated Reactome terms in EGFR^ΔEgr2^ JAK1/2i anti-IL-17A compared to EGFR^ΔEgr2^ JAK1/2i. **(F)** Gene expression within mouse groups of positively-correlated genes between human EGFRi-treated patients and EGFR^ΔEgr2^ mice from Fig. 1C.

Taken together, adjunctive IL-17A blockade did not compromise the therapeutic benefits of JAK1/2 inhibition but instead provided additional benefits by reducing antigen presentation in KCs and attenuating the inflammatory cutaneous environment. These findings identify IL-17A as a contributor to residual inflammatory pathology and demonstrate that selectively targeting distinct immune pathways can further promote tissue homeostasis following JAK1/2 inhibition.

## Discussion

Our work identified targeted impacts of therapeutic JAK1/2 inhibition that reshape a multifaceted inflammatory environment to enable tissue regeneration. At single-cell resolution, we demonstrated how redirection of the immune response promotes resolution of chronic inflammation and confirmed therapeutic success, as evidenced by barrier restoration, repopulation of diverse hair follicle cell types and hair regrowth.

Previously, we observed that despite an almost complete loss of CD34^+^ bulge stem cells, Sox9^+^ stem cells were able to regenerate the hair follicle in progressive scarring alopecia, provided that topical JAK1/2i was administered in a timely manner.^7^ We now investigated the rudimentary hair follicle cell populations before and during JAK1/2i treatment, to provide an in-depth profile of potential regenerative targets. The major HF cluster in the degenerated mutant condition was the *OB_Camk1d* cluster, defined by upregulation of *Camk1d*, encoding calcium/calmodulin-dependent protein kinase 1D (CAMK1D). Interestingly, CAMK1D was previously shown to mediate tumor cell resistance to cytotoxic T lymphocyte killing by inhibiting apoptosis.^24^ FasL exposure of tumor cells *in vitro* triggered a rise in intracellular Ca^2+^, activating CAMK1D, which then in turn suppressed caspases-3, –6, and –7. *Camk1d* upregulation could therefore constitute a survival mechanism against a prolonged immune attack to establish a more resistant niche from where the HF can be regenerated once the inflammation subsides. Notably, the *OB_Camk1d* cluster exhibited a high proportion of unspliced transcripts compared to all other clusters. This reflects either a high transcriptional activation state or a disruption in splicing processes, which are often associated with cell stress, immune signaling and altered differentiation dynamics.^25^

Upon JAK1/2 inhibition we observed a diverse recovery of different hair follicle cell populations. Most strikingly, the upper HF region was reinstated, a region crucial for antimicrobial defense and barrier function.^26^ Notably, JAK1/2 inhibition restored key tight and adherens junction proteins, such as claudin-1, in the upper hair follicle as early as three days post therapy start, marking it the most prominent immediate cutaneous benefit. Restored cutaneous barrier integrity restricts both type 2 immunity and microbial exposure.^27^ Overall, this indicates a potential direct effect of JAK1/2 inhibition on barrier repair and explains its promising therapeutic potential in AD, which is driven by barrier defects and type 2 immunity.

In line with these therapeutic observations on the organism level, among the T cells, CD8 followed by CD4 αβT cell compartments show the strongest JAK-STAT pathway activity and, consequently, the strongest deactivation upon JAK1/2 inhibition. On the other hand, the type 3 response carried out by the γδTC populations are not JAK-STAT-dependent and remained active upon initial JAK1/2 inhibition. Surprisingly, during the initial and the transitional therapeutic phases, both the CD8 and CD4 αβT cell compartments switched from their respective type 1 and 2 immunity to type 3 signatures with IL-17A expression as a hallmark.

Notably, IL-17A, together with other JAK-STAT-independent cytokines such as TNFα and IL-1β, is essential for antifungal and antimicrobial immunity.^28^ This is largely due to their role in promoting neutrophil recruitment and production of antimicrobial peptides, particularly relevant for *S. aureus* clearance.^29^ In line with this, we observed a phenotypic shift in neutrophils under JAK1/2i. Reflecting changes seen in lymphocytes, neutrophils displayed a switch from *Ifngr1* and *Il4ra* to *Il17ra* expression. This happened alongside an upregulation of genes associated with antimicrobial defense including *Lcn2*, *S100a8* and *S100a9*.^30^ In earlier studies, we observed that neutrophils numbers decreased following antibiotic treatment in mice with epidermal EGFR deletion.^14^ However, neutrophil numbers did not change or even increased relative to other immune cells in the resolution phase (4 weeks) of JAK1/2i treatment.^7^ Therefore, JAK1/2i is capable of boosting certain immune responses, possibly by removing interfering signals.

In the skin, IL-17 promotes keratinocyte proliferation, which is beneficial in wound healing processes, however, excessive activation of the IL-17 pathway can also drive pathological responses, as seen in psoriasis.^31^ The relevance of IL-17 and type 3 immunity is also increasingly recognized in AD, although type 2 immunity is still considered the primary driver.^32^ An elevated Th17 pathway has been reported in several AD subtypes, including pediatric AD, older patients, and patients of Asian ethnicity, where it is thought to contribute to epidermal hyperplasia and lichenification.^33^ Lesional skin, which is commonly colonized by pathogenic bacterial has also been associated with an enrichment of additional Th17 and also Th1 responses.^34–36^ Multifaceted immune responses are also present in our mouse model, in which type 2 signals (i.e. TSLP) dominate the early disease phase following initial skin barrier disruption during hair eruption.^14,15^ Only in later phases, type 1 and type 3 responses emerge, whereas the latter are driven by bacterial dysbiosis.^16,14^

Our data from the early therapeutic phase indicate that JAK1/2 inhibitors do not affect IL-17A signaling, raising the possibility that additional IL-17A blockade may provide further benefit in skin conditions with IL-17-supported pathology. In support of this, concomitant blockade of IL-17A and JAK1/2 in our mouse model, resulted in further reduction of antigen presentation as seen by MHC-II down-regulation in HF and IFE keratinocytes, while preserving improvements in barrier integrity and microbial control. Considering the potent antimicrobial and antifungal role of type 3 immunity, excessive IL-17A inhibition could lead to detrimental effects in patients with skin barrier defects.^21,31,37^ However, such adverse effects were not observed. This suggests that IL-17A modulation in combination with JAK1/2 inhibition represents a promising therapeutic strategy to address the limitations of JAK inhibitor monotherapy and provides a rationale for exploring combination therapies with biologics.

In conclusion, our findings demonstrate that JAK1/2i does not simply induce broad immunosuppression but instead selectively attenuates pathologic type 1– and type 2-driven pathways, linked to cytotoxicity and barrier dysfunction, while preserving or enhancing type 3-mediated antimicrobial responses and barrier restoration. Importantly, our preclinical data support a model in which immune recalibration, rather than global suppression, underlies the clinical efficacy of JAK inhibitors in chronic inflammatory conditions and that additionally inhibiting selected pathologic cytokines like IL-17A can enhance therapeutic efficiency. Defining the contexts in which these strategies are most effective will be essential to guide precision treatment, identify predictors of therapeutic response, and inform rational combination approaches.

## Materials and methods

### Mice

EGFR^ΔEgr2^ mice were generated as previously described^7^ by crossing EGFR^fl/fl^ mice from Natarajan, Wagner and Sibilia^38^ with Egr2^Cre^ mice (The Jackson Laboratory, strain #025744). All mice were bred and housed in the facilities of the Medical University of Vienna. Animal experiments were approved by the Animal Experimental Ethics Committee of the Medical University of Vienna and the Austrian Federal Ministry of Science and Research (animal license number: GZ BMWFW-66.009/0319-V/3b/2019). Mice were randomly picked for experiments and sexes equally allocated, including all relevant controls as in treated and untreated wild-type littermates of the EGFR^ΔEgr2^ mice.

### Topical JAK inhibitor treatment

Mice were treated topically with the JAK1/2i ruxolitinib as previously described^7^. Briefly, ruxolitinib was dissolved in DMSO at a concentration of 3%. Ruxolitinib or vehicle (DMSO) was applied topically by spreading a thin layer on back skin, ear skin and tail skin. For adult mice, a volume of about 120 µl was used depending on the size of the mouse. This was repeated daily for the durations stated in figure legends.

### IL-17A blockade

To inhibit IL-17A, mice were injected intraperitoneally with 200 µg anti-mouse IL-17A (Clone: 17F3, InVivoMab) or mouse IgG1 isotype control (Clone: MOPC-21) every two days for a total duration of two weeks.

### TEWL measurement

According to the manufacturer’s recommendations, the transepidermal water loss (TEWL) was measured with a Tewameter® TM 300 probe connected to the MDD4-display device (Courage + Khazaka).

### Immunohistochemistry

For immunofluorescence stainings of back skin, formalin-fixed paraffin-embedded tissue was cut into 4 µm thin sections. Sections were deparaffinized and rehydrated, followed by antigen retrieval in DAKO antigen retrieval solution (pH=9) for 15 min using a 2100 retriever cooker. After cooling to room temperature, tissues were circled with a PAP pen and washed with PBS. Tissues were blocked for 1 h at room temperature in 5% horse serum and 0.1% Triton-x-100 in PBS. Followed by incubation with primary antibodies at 4°C overnight or 2 h at room temperature. Sections were washed with PBS, and incubated for 2 h with secondary antibodies and Hoechst or DAPI. After a final wash with PBS, slides were mounted. To keep sections from drying out and protect them from light, slides were stored in a dark humidified slide chamber for the whole staining process. Images were taken using an Olympus iX83 WF Automated Widefield FL or Olympus/Evident VS200 BF+FL+Pol Slidescanner and analyzed using the CellSens software. Quantification of claudin-1^+^ area in hair follicle cell membrane was carried out using HALO AI (Indica Labs) software.

### Fluorescence-Activated Cell Sorting

To make epidermal single-cell suspensions, ear and back skin were processed as follows. For back skin, subcutaneous fat was gently scraped off and skin was laid dermis-side down on 0.8% trypsin in PBS for 45 min at 37°C. Ears were split in dorsal and ventral sides and floated dermis-side down on 0.8% trypsin in PBS for 30 min at 37°C. After trypsin-incubation, the epidermal layer was separated from the dermis and further processed in 250 µg/ml DNase-I in PBS by 30 min at 37°C in a shaker set to 650 rpm. The resulting mixture was filtered using a 70 µm cell strainer. To achieve a single-cell suspension, skin-draining lymph nodes were mechanically homogenized and washed through a 70 μm cell strainer with PBS 2% FBS.

### Intracellular cytokine staining

Single-cell suspensions of back skin and lymph nodes were incubated for 4 h at 37°C in 24-well cell culture plates in 500 µl complete RPMI (Sigma) with cell stimulation cocktail (eBioscience TM 00-4975) or protein transport inhibitor cocktail (eBioscience, 00-4980) for the negative control. After cell stimulation, cells were resuspended and washed, followed by extracellular staining as described below and fixation with IC fixation Buffer (eBioscience, 00-8222). Intracellular cytokine staining was carried out according to manufacturer recommendations using eBioscience intracellular fixation & permeabilization buffer set (Cat. Nr. 88-8824).

### Extracellular staining

Single-cell suspensions were stained as previously described.^7^ Briefly, cell suspensions were incubated with anti-CD16/32 Fc block (Biolegend). Dead cells were identified using SYTOX™ Blue Dead Cell Stain (1:1000, Thermo Fisher Scientific) or Zombie Aqua (1:500, Biolegend) for samples that were subsequently fixed. Extracellular antibody staining (see table) incubated for 30 min at 4°C. Cells were analyzed using an LSR-II flow cytometer (BD Biosciences) and FlowJo Software.

### Single-cell RNA sequencing

For scRNA-seq, 60.000 live CD45^+^ immune cells and CD45^-^Sca-1^-^hair follicle cells were sorted from epidermal single-cell solutions without influencing cell type numbers using a BD FACSMelody™ Cell Sorter (Biosciences). Single-cell cDNA libraries were generated using the droplet-based Chromium Next GEM Single Cell 5’ Kit v2 (Cat.nr. PN-1000265, 10x Genomics) following manufacturer’s instructions.

### scRNA-seq analysis

Preprocessing of the sequencing data was performed using the CellRanger pipeline version 7.1.0 using the mouse transcriptome mm10-2020-A and with the option to include introns. CellRanger output files were then processed with various R packages. SoupX^39^ (v1.6.2) was used to correct for ambient RNA contamination. DropletUtils (v1.20.0) was used for conversion of data objects into appropriate formats. Doublet exclusion was performed with scDblFinder^40^ (v1.14.0). For downstream analysis, Seurat version 4.3.0 and up^41,42^ was employed. All samples were QCed individually before merging samples into one Seurat object. Merging was followed by clustering and manual cell type annotation based on marker genes. Markers were determined using Seurat’s FindConservedMarkers function (parameters different from default: only.pos = TRUE). Larger clusters were first downsampled to 100 cells to improve speed. First-level clusters were grouped into main classes: HF, Lymphoid, DETC and Myeloid. These were subsetted and analyzed separately. Each main class went through further QC, removing of low-quality cells, clustering and annotation. For the main dataset shown in Fig. 1, cleaned-up main classes were merged together again to ensure that only high-quality cells were included. Comparisons between conditions within clusters were performed using the FindMarkers function with default parameters. Cell-cell interactions were determined using CellChat^43^ (v2.2.0). CellChat was run according to the tutorials describing the analysis of single-cell transcriptomics data provided in the GitHub repository (github.com/jinworks/CellChat). The min.cells parameter in the filterCommunication function was set to 3. To infer signaling activity using PROGENy, the progeny^44^ package (v1.26.0) was used. Visualization and plotting were aided by SeuratExtend (v1.0.8) for marker dotplots and barplots, scCustomize (v3.0.1) dotplots for T cell polarization^45^, pheatmap (v1.0.12) for heatmaps, dittoSeq (v1.20.0) also for heatmaps.^46^

### RNA velocity

RNA velocity analysis on hair follicle cells was performed via the tool scVelo^47^ (v0.3.3) using Python (v3.10). Cells labelled as non-keratinocytes were excluded from the RNA velocity analysis. Spliced and unspliced transcript counts were obtained from velocyto^25^ (v0.17.17) using the 10X Chromium command with default parameters and stored as loom-files. The required RepeatMasker file was downloaded from the UCSC table browser.^48^ Gene annotations (Release M25, GRCm38.p6) were downloaded from Gencode.^49^ The Seurat object containing the scRNA-seq data was converted to an AnnData object via SeuratDisk. The information on spliced and unspliced transcript counts was added to the AnnData object via Scanpy (v1.9.8) and scVelo. The RNA velocity was computed with scVelo (‘dynamical’ mode) using the standard workflow and default parameters. Untransformed (raw) scRNA-seq were used as input to scVelo. The computed velocity vector field was projected on the UMAP embedding. Besides the analysis on the combined set of conditions (wt_dmso, wt_ruxo, egr2_dmso, egr2_ruxo), the RNA velocity was also computed separately for each condition by subsetting the AnnData object prior to the scVelo analysis. These separate analyses are shown in Fig. 2 and S2. The unspliced/spliced ratio shown in Fig. 2D was computed per cell from raw velocyto counts as log₂((unspliced+1)/(spliced+1)). Pairwise differences between conditions were assessed with two-sided Mann–Whitney U tests, restricted to the four comparisons in which a single factor varied (treatment within each genotype and genotype within each treatment). P-values were adjusted across these four tests by the Benjamini– Hochberg procedure and are reported as q-values.

### Bacterial skin swabs

For measuring the bacterial load of WT and EGFR^ΔEgr2^ mice before and after 2-4 weeks JAK1/2i treatment, all conditions were housed separately to avoid transfer of bacteria between groups. Backskin was swabbed 30 times with gentle pressure using sterile cotton swabs that were pre-wetted in sterile PBS. Cotton swab tips were then submerged in 1.5 ml Eppendorf tubes containing 1 ml sterile PBS, cut off from the stick and left to incubate after 5 shakes of the tube. 10-20 µl PBS containing swab tips was plated on COS Columbia + 5% Sheep Blood agar plates by BioMérieux (cat. 43041).

### Public dataset acquisition and re-analysis

Bulk RNA-sequencing data from the public patient dataset published by Rutkowski *et al.*^20^ was retrieved from the European Nucleotide Archive (ENA, accessible under: https://www.ebi.ac.uk/ena/browser/view/ERP152609) using iSeq^50^ for secondary bioinformatic re-analysis. Raw sequencing FASTQ files were processed using the nf-core/rnaseq (v3.26.0)^51,52^ workflow using Nextflow (v26.04.6).^53^ Transcript-level abundances were quantified in pseudo-alignment mode using Salmon. Transcript counts and length estimates were aggregated to gene-level matrices using tximport. These matrices were directly passed to the nf-core/differentialabundance (v.2.0.0)^54^ workflow for evaluation of differential gene expression was evaluated using DESeq2 (v.1.34.0).^55^ Gene Ontology (GO) gene set enrichment analysis (GSEA) was performed using clusterProfiler (v4.18.4).^56,57^ For visualization (including generation of Semantic Space plot, cnetplot, and enrichment plot) the accompanying package enrichplot (v.1.30.5)^58^ was used. ComplexHeatmap (v.2.26.1) R package was used for generating heatmaps.

### Bulk mRNA sequencing of mouse skin

Full-thickness back skin biopsies from WT and EGFR^ΔEgr2^ mice were collected and stored in RNAlater (Sigma-Aldrich, cat. R0901) at –80°C. For RNA extraction, tissue was allowed to thaw and 3 mm rounds were taken using 3 mm biopsy punches (KAI Medical, cat. BP-30F). Remaining RNAlater was removed using a tissue before proceeding with the RNeasy® Fibrous Tissue Mini Kit (QIAGEN, cat. 74704). Following the kit’s instructions, RLT buffer was supplemented with 20 µl 2 M dithiothreitol (DTT) per 1 ml RLT buffer. For the tissue homogenization step, skin pieces were added to Precellys CK28 bead tubes (ref. P000911-LYSK1-A.0) together with 300 µl RLT buffer with DTT. Tissues were then processed in a Precellys Evolution Touch homogenizer by Bertin Technologies set to 3 cycles of 6000 rpm for 20 s each, with a 10 s pause between cycles. Lysates were then further processed by following the RNeasy® Fibrous Tissue Mini Handbook.

Purified RNA was sent to Novogene Europe (Munich) for strand-specific mRNA sequencing library preparation. Libraries were sequenced on an Illumina platform with a target depth of 30 million reads per sample.

### Analysis of mouse bulk mRNA sequencing

Basic bioinformatics analysis was provided by Novogene. Briefly, Quality filtering of raw FASTQ reads was performed using fastp software to remove adapter sequences and low-quality bases. Clean paired-end reads were mapped to the Mus musculus reference genome GRCm39 using HISAT2 (v2.2.1). Gene-level read counts were generated using featureCounts (v2.1.1). Differential expression analysis was performed using the DESeq2 R package (v1.42.0).^55^ Genes with a P-value ≤ 0.05 and absolute log2 fold change ≥ 1.0 were defined as significantly differentially expressed for downstream functional analyses. Reactome pathway enrichment analyses were implemented using clusterProfiler (v4.8.1).^56,57^ For the comparison of mouse and human patient gene expression, the log2 fold changes of WT vs. EGFR^ΔEgr2^ and healthy control vs. patient were plotted by overlapping orthologous DEGs (adjusted p-value ≤ 0.05, absolute log2 fold change ≥ 0.5). Ortholog mapping was performed using the *gorth* function of the gprofiler2 (v. 0.2.4) R package. The correlation dot plot and heatmaps of correlated genes were visualized using ggplot2 (v.4.0.3) and ComplexHeatmap (v.2.26.1) R packages.

### Statistics

Statistical analyses for quantitative data of TEWL, immunofluorescence microscopy, bacterial abundance and FACS were performed in GraphPad Prism 8. The specific statistical tests used and the number of biological replicates per group are noted in the respective figure legends. P-values above 0.05 were omitted from graphs for better readability. For bioinformatics analyses including DESeq2 and GSEA, the Benjamini Hochberg method was used for determining the adjusted p-value.

## Acknowledgements

This work was supported by grants from the Austrian Science Fund (FWF), grant number DOC188 and PAT5056523 to T.B. R.J. was supported by the Austrian Academy of Sciences (ÖAW, Dimitrov-Legat scholarship). This work was further supported by the following grants acquired by MS: FWF DocFunds DOC32-B28 and DOC188 and the European Research Council (ERC) Advanced grant (ERC-2015 AdG TNT-Tumors 694883).

We thank Dana Krauss and Thomas Mohr for advice on bioinformatical analysis methods; Flora Tomaselli and Veronica Hofer for technical assistance; Martina Hammer for help with maintaining the mouse colonies; and the FACS, imaging and histology facilities of the Center for Cancer Research for their excellent technical assistance.

## Author contributions (using CRediT Taxonomy)

Regina Jin: Conceptualization; Data curation; Validation; Investigation; Formal analysis; Visualization; Methodology; Writing – original draft; Writing – review and editing.

Karoline Strobl: Conceptualization; Data curation; Investigation; Validation; Formal analysis; Visualization; Methodology; Writing – original draft; Writing – review and editing.

Lena Artner-Gent: Data curation; Investigation; Validation; Formal analysis. Johannes Tüchler: Formal analysis; Methodology; Visualization.

Marina Gladkova: Investigation; Writing – review and editing. Ulrike Mann: Data curation; Investigation.

Matthias Farlik: Resources; Supervision. Iros Barozzi: Formal analysis; Supervision.

Maria Sibilia: Resources; Review and editing; Supervision; Funding acquisition; Project administration.

Thomas Bauer: Conceptualization; Visualization; Resources; Supervision; Funding acquisition; Project administration; Writing – original draft; Writing – review and editing.

## Data availability

The scRNA-seq dataset will be made available through the Gene Expression Omnibus (GEO) upon publication.

