## Supplementary Figures for "JAK1/2 Inhibition Restores Tissue Integrity by Rewiring Type 1/2 Immunity Toward Targetable IL-17 Responses"

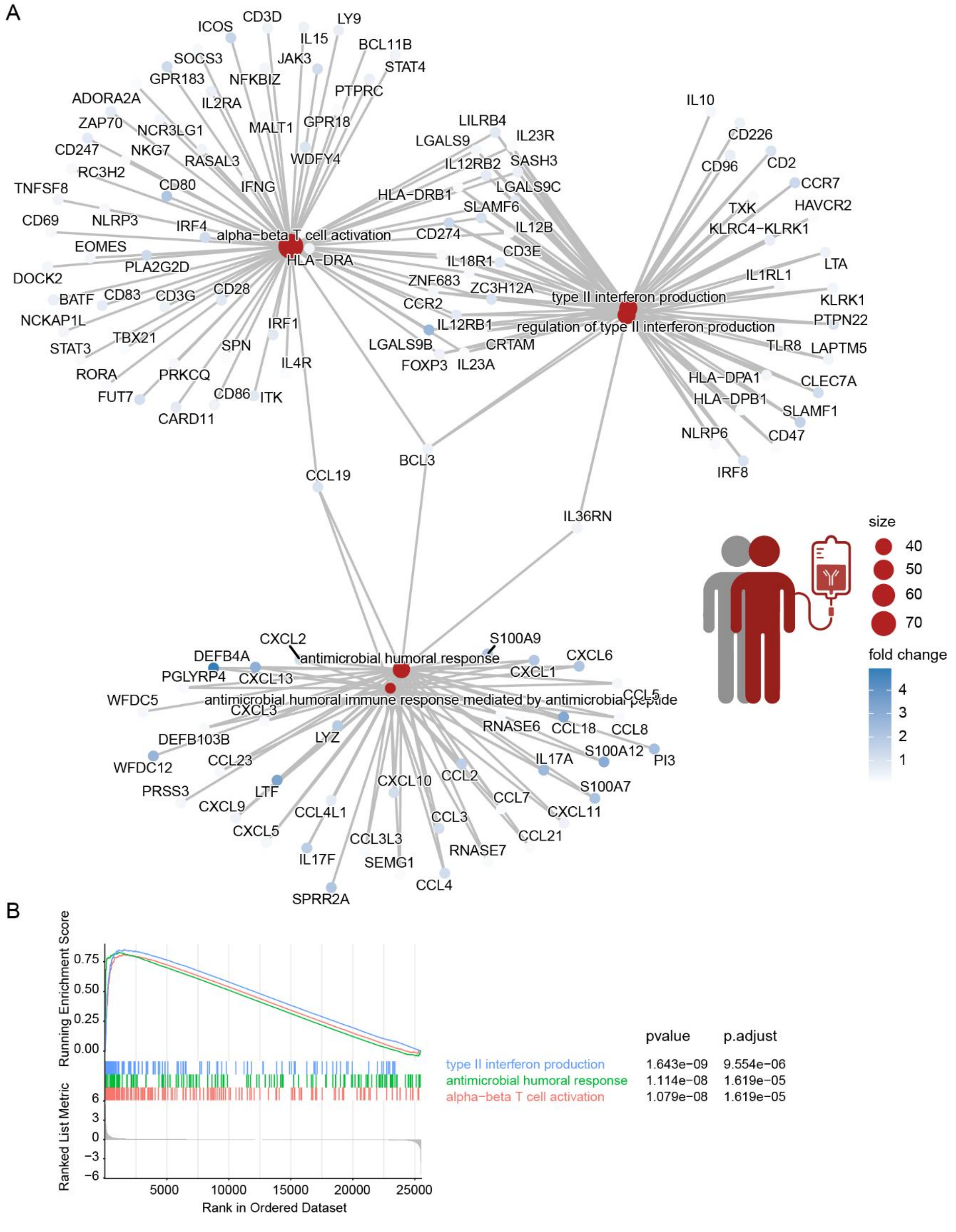


**Fig. S1. Multifaceted inflammatory landscape associated with long-term EGFR inhibition.**

GSEA was performed between EGFRi-treated patients and healthy control scalp skin. **(A)** ClusterProfiler cnetplot of top 5 enriched genesets. **(B)** Running geneset enrichment plot of top 3 enriched genesets.

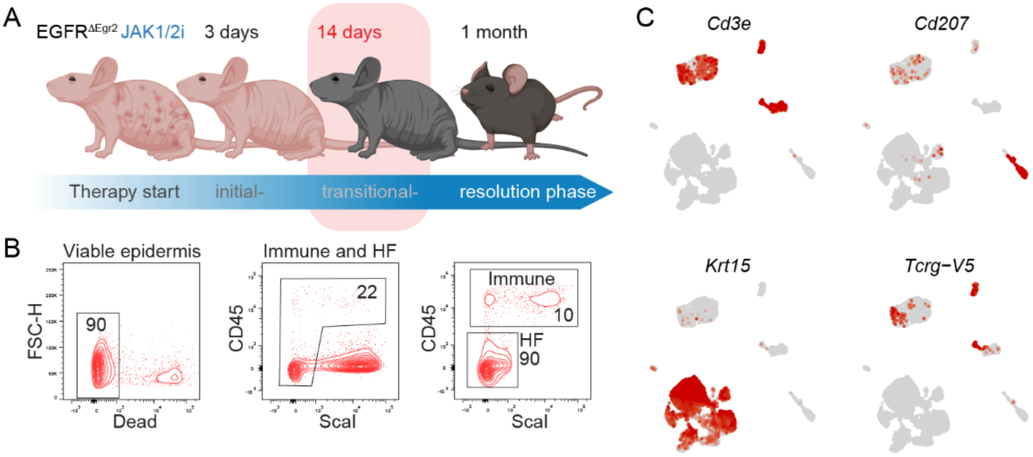


**Fig. S2. Transitional therapeutic stage of JAK1/2 inhibition in a mouse model of chronic skin inflammation.**

**(A)** Schematic overview of therapeutic stages. **(B)** Gating strategy used for sorting cells for the scRNA-seq and FACS plots shown in Fig. 1G. **(C)** Relative expression of marker genes for main classes of the scRNAseq dataset, using UMAP embedding of single cells.

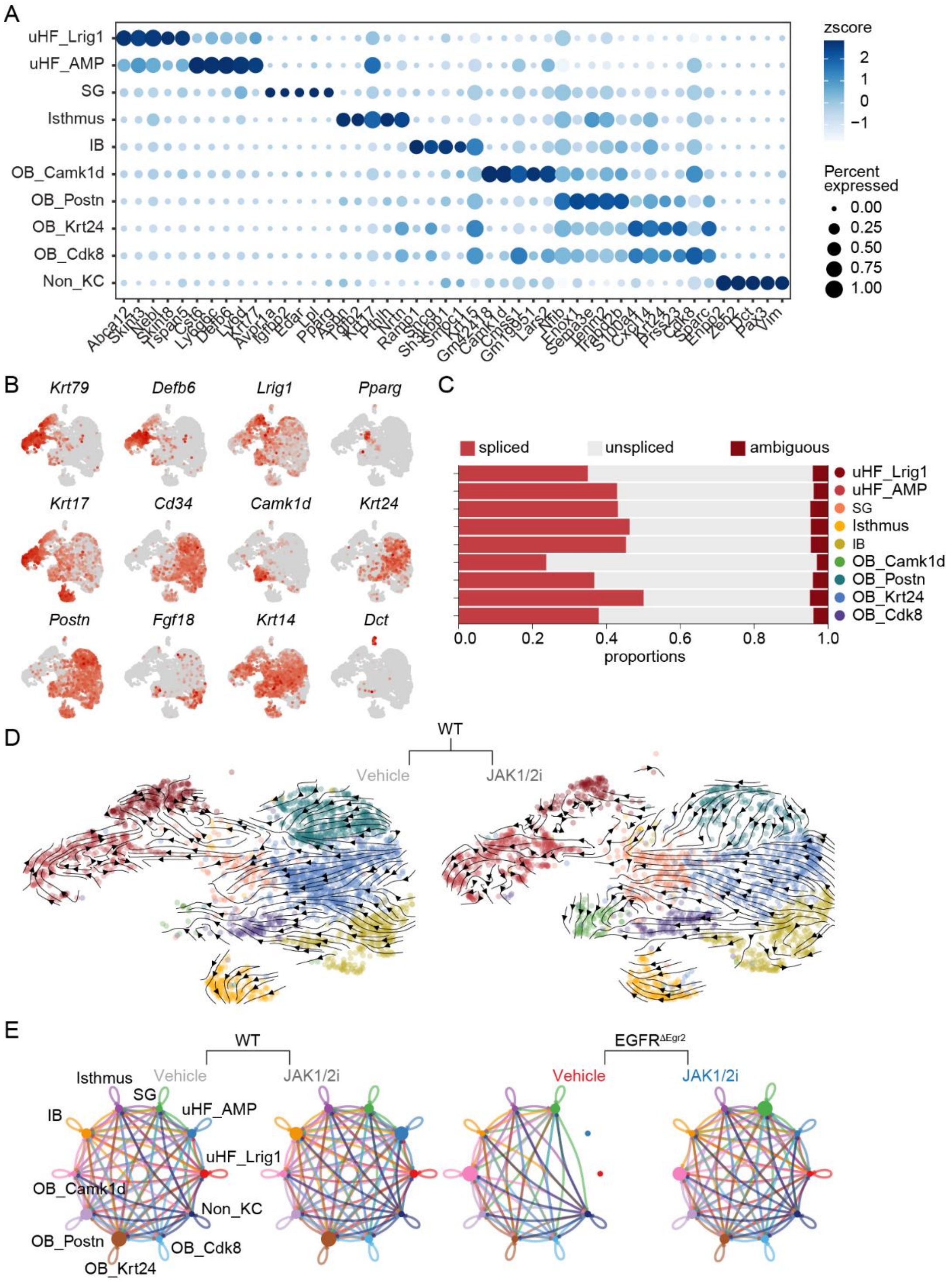


**Fig. S3. Hair follicle cell populations during inflammation and JAK1/2i therapy**.

**(A)** Average expression of top 5 marker genes for each HF subset depicted in a dotplot. **(B)** UMAP of HF re-clustering, colored by expression of key marker genes. **(C)** Proportion of spliced and unspliced transcripts in each cluster across conditions, determined by velocyto. **(D)** RNA velocities of WT conditions projected onto UMAP embeddings and visualized as streamlines. **(E)** Circle plot depicting total number of cell-cell interactions as determined by CellChat.

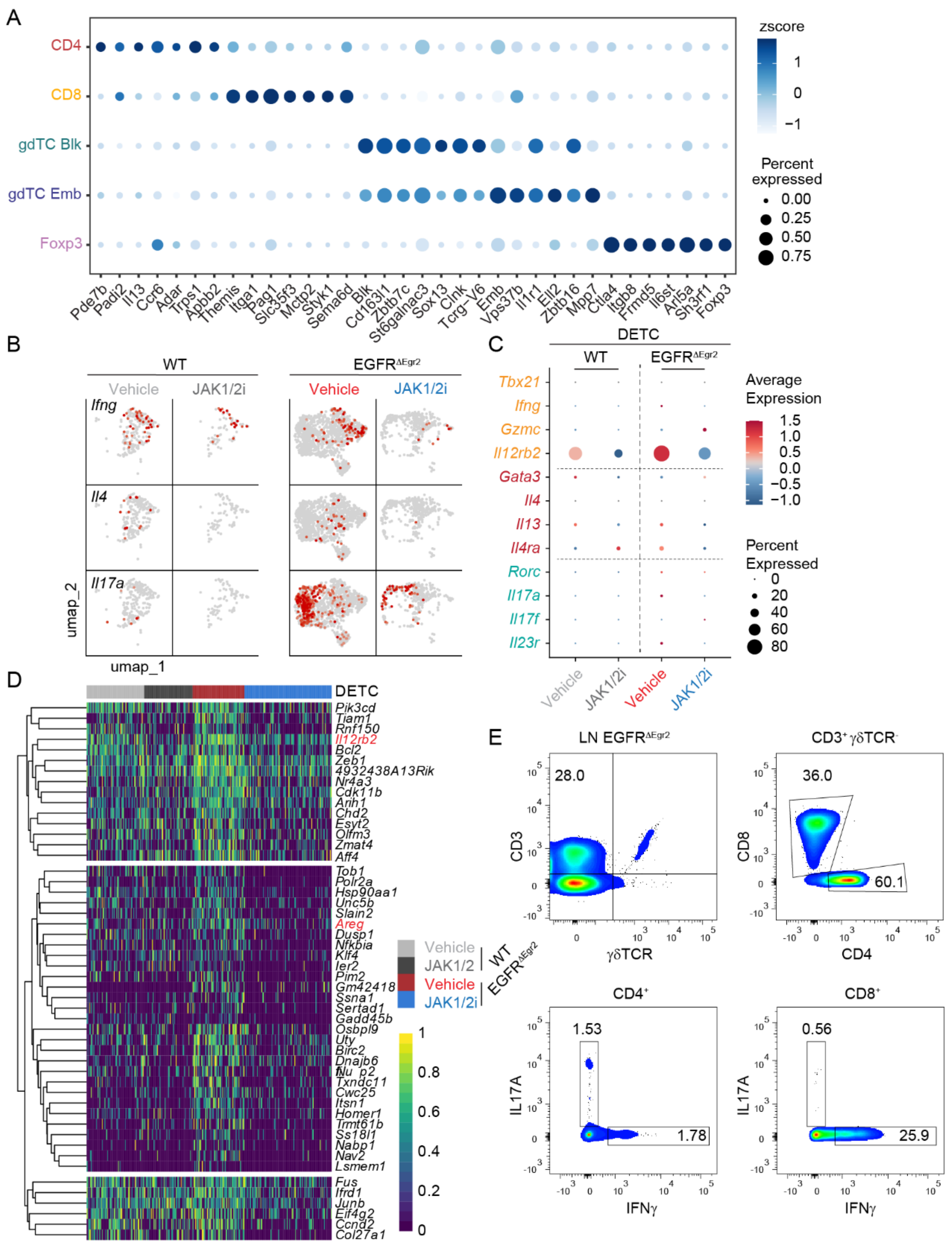


**Fig. S4. Epidermal T cell subsets following JAK1/2 inhibition.**

**(A)** Average expression of top 7 marker genes of each Lymphoid subset depicted in a dotplot. **(B)** *Ifng*, *Il4* and *Il17a* expression plotted on UMAP of each condition. **(C)** Expression of T cell polarization genes in DETCs. **(D)** Heatmap depicting gene expression in DETCs, showing expression of Top 50 most significant downregulated DEGs in between JAK1/2i-treated EGFR$^{\Delta Egr2}$ compared to vehicle-treated EGFR$^{\Delta Egr2}$. **(E)** Gating strategy for intracellular cytokine FACS analysis for Fig. 4F.

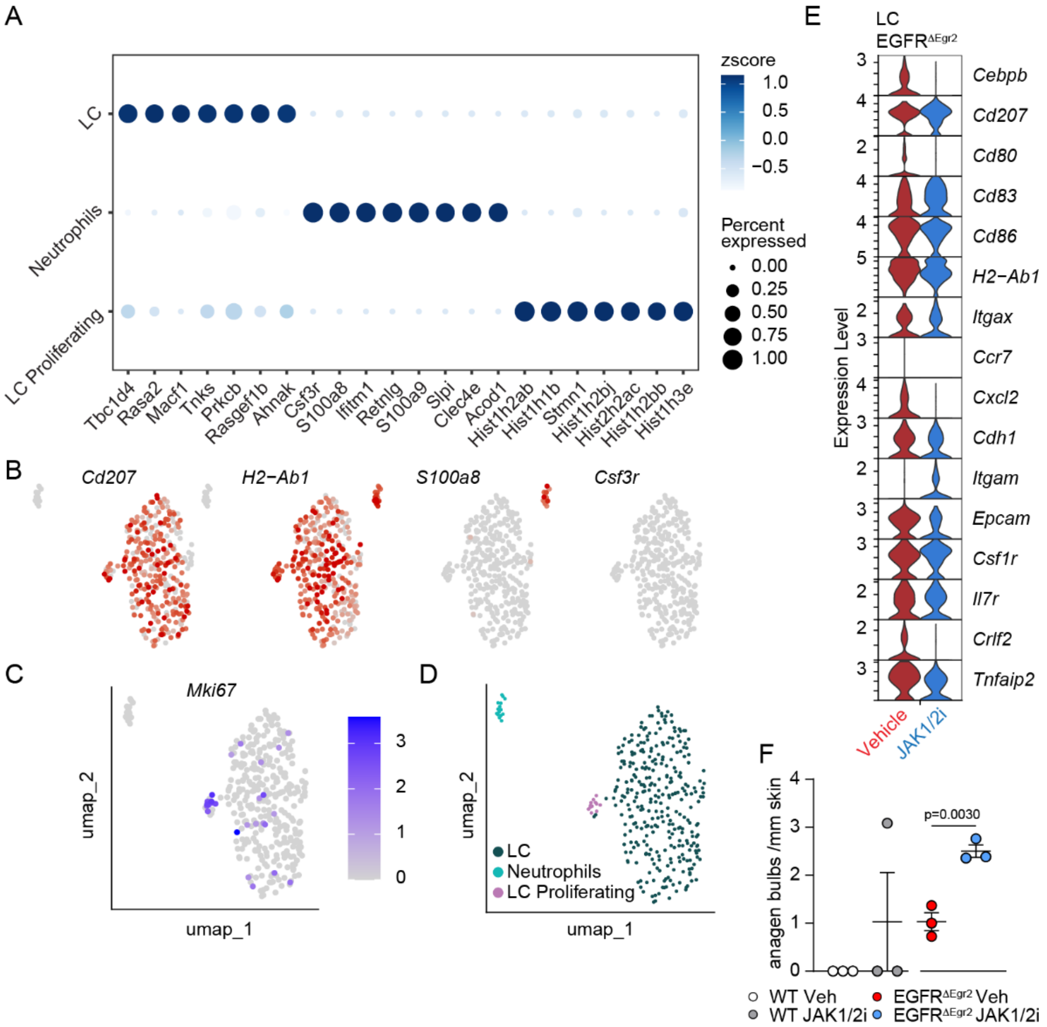


**Fig. S5. Analysis of Myeloid subsets during JAK1/2i therapy response.**

**(A)** Dotplot showing expression of Top 7 significant marker genes between Myeloid clusters found by Seurat's FindMarkers function. **(B)** Marker gene expression projected onto UMAP. **(C)** Mki67 expression in Myeloid cells**. (D)** UMAP representation of Myeloid subsets. **(E)** Violin plot depicting relative expression of selected genes in LC subset of $EGFR^{\Delta Egr2}$ mice treated with vehicle or JAK1/2i for 2 weeks. **(F)** Quantification of anagen hair bulbs from back skin sections from vehicle- or JAK1/2i-treated WT and $EGFR^{\Delta Egr2}$ mice (n ≥ 3). t-tests were used to compare vehicle- and JAK1/2i-treated mice.

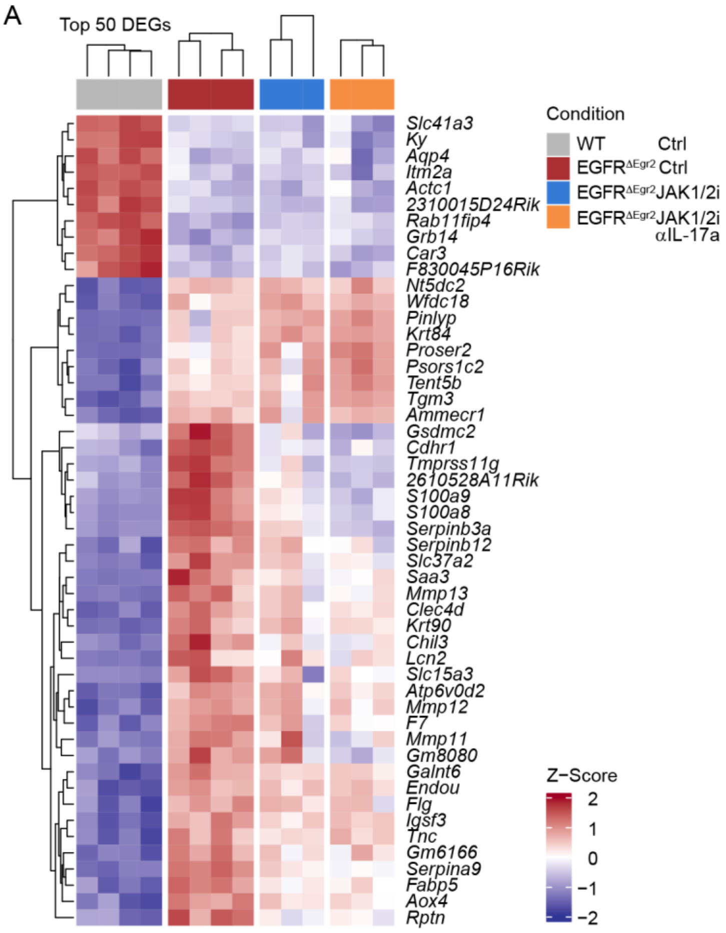


**Fig. S6. Bulk RNAseq profiling of EGFR$^{\Delta Egr2}$ mice treated with JAK1/2i alone or in combination with IL-17A blockade.**

**(A)** Heatmap showing the top 50 significantly differentially expressed genes between WT and EGFR$^{\Delta Egr2}$ mice.
